# Integrative Network Fusion: a multi-omics approach in molecular profiling

**DOI:** 10.1101/2020.04.01.020685

**Authors:** Marco Chierici, Nicole Bussola, Alessia Marcolini, Margherita Francescatto, Alessandro Zandonà, Lucia Trastulla, Claudio Agostinelli, Giuseppe Jurman, Cesare Furlanello

## Abstract

Recent technological advances and international efforts, such as The Cancer Genome Atlas (TCGA), have made available several pan-cancer datasets encompassing multiple omics layers with detailed clinical information in large collection of samples. The need has thus arisen for the development of computational methods aimed at improving cancer subtyping and biomarker identification from multi-modal data. Here we apply the Integrative Network Fusion (INF) pipeline, which combines multiple omics layers exploiting Similarity Network Fusion (SNF) within a machine learning predictive framework. INF includes a feature ranking scheme (rSNF) on SNF-integrated features, used by a classifier over juxtaposed multi-omics features (juXT). In particular, we show instances of INF implementing Random Forest (RF) and linear Support Vector Machine (LSVM) as the classifier, and two baseline RF and LSVM models are also trained on juXT. A compact RF model, called rSNFi, trained on the intersection of top-ranked biomarkers from the two approaches juXT and rSNF is finally derived. All the classifiers are run in a 10×5-fold cross-validation schema to warrant reproducibility, following the guidelines for an unbiased Data Analysis Plan by the US FDA-led initiatives MAQC/SEQC. INF is demonstrated on four classification tasks on three multi-modal TCGA oncogenomics datasets. Gene expression, protein abundances and copy number variants are used to predict estrogen receptor status (BRCA-ER, N=381) and breast invasive carcinoma subtypes (BRCA-subtypes, N=305), while gene expression, miRNA expression and methylation data is used as predictor layers for acute myeloid leukemia and renal clear cell carcinoma survival (AML-OS, N=157; KIRC-OS, N=181). In test, INF achieved similar Matthews Correlation Coefficient (MCC) values and 97% to 83% smaller feature sizes (FS), compared with juXT for BRCA-ER (MCC: 0.83 vs 0.80; FS: 56 vs 1801) and BRCA-subtypes (0.84 vs 0.80; 302 vs 1801), improving KIRC-OS performance (0.38 vs 0.31; 111 vs 2319). INF predictions are generally more accurate in test than one-dimensional omics models, with smaller signatures too, where transcriptomics consistently play the leading role. Overall, the INF framework effectively integrates multiple data levels in oncogenomics classification tasks, improving over the performance of single layers alone and naive juxtaposition, and provides compact signature sizes^1^.

## 1 INTRODUCTION

The challenge of integrating multi-omics data is as old as bioinformatics itself (1, 2), but, despite the wide literature, it remains an open issue nowadays, even worth being funded by major institutions^2^.

Our study introduces Integrative Network Fusion (INF), a reproducible network-based framework for high-throughput omics data integration that leverages machine learning models to extract multi-omics predictive biomarkers. Originally conceptualized and tested on multi-omics metagenomics data in an early preliminary version (3, 4), INF combines the signatures retrieved from both the early-integration approach of variable juxtaposition (juXT) and an intermediate-integration approach (SNF (5)), to find the optimal set of predictive features. In particular, first a set of top-ranked features is extracted by juXT by a classifier, here Random Forest (RF) and linear Support Vector Machine (LSVM). Then, a feature ranking scheme (rSNF) is computed on SNF-integrated features and finally a RF model (rSNFi) is trained on the intersection of two set of top-ranked features from juXT and rSNF, obtaining an approach that effectively integrates multiple omics layers and provides compact predictive signatures. Selection bias and data-leakage effects are controlled by performing the experiments within a rigorous Data Analysis Plan (DAP) to warrant reproducibility, following the guidelines of the US FDA-led initiatives MAQC/SEQC (6, 7, 8). In particular, to alleviate the computational burden of the full DAP pipeline, an approximating DAP is designed to lighten computing without significantly affecting the results. Further, experiments are run on samples with randomly shuffled labels as a sanity check versus overfitting effects and, finally, INF robustness is verified by testing on different train/test splits.

We test INF on three datasets retrieved from the TCGA repository, to predict either the estrogen receptor status (ER) or the cancer subtype on the breast invasive carcinoma (BRCA) dataset, and to predict the overall survival (OS) on the kidney renal clear cell carcinoma (KIRC) and acute myeloid leukemia (AML) datasets. Overall, INF improves over the performance of single layers and naive juxtaposition on all four oncogenomics tasks, extracting a biologically meaningful compact set of predictive biomarkers. Notably, the transcriptomics layer is prevalent inside the inferred INF signatures, consistently with published findings (9).

The INF framework is currently designed to integrate an arbitrary number of one-dimensional omics layers. We plan to further extend the framework by enabling the integration of histopathological features extracted from whole slide images (10) or deep features from radiological images (11) extracted by deep neural network architectures, carefully addressing all potential caveats (12).

To further foster reproducibility and support users and future developers, the full code of this benchmark is publicly shared on the GitLab repository https://gitlab.fbk.eu/MPBA/INF. Additional information is included in the Supplementary Material available on the publisher’s website, while the full set of experimental data can be accessed at http://dx.doi.org/10.6084/m9.figshare.12052995.v1.

## 2 MATERIALS AND METHODS

### 2.1 Data

Three multi-modal cancer datasets generated by The Cancer Genome Atlas (TCGA) Research Network (https://www.cancer.gov/tcga) and four classification tasks are considered in this study. Protein abundance (*prot*), gene expression (*gene*) and copy number variants (*cnv*) are used to predict breast invasive carcinoma (BRCA) estrogen receptor status (0: negative; 1: positive) and subtypes (luminal A, luminal B, basal-like, HER2-enriched). Methylation (*meth*), gene expression (*gene*) and microRNA expression (*mirna*) are used to predict acute myeloid leukemia (AML) and kidney renal clear cell carcinoma (KIRC) overall survival (0: alive; 1: deceased). The number of samples and features for each omic layer and classification task are detailed in Table 1; class balance, split by dataset, is reported in Table 2.

**Table 1.**
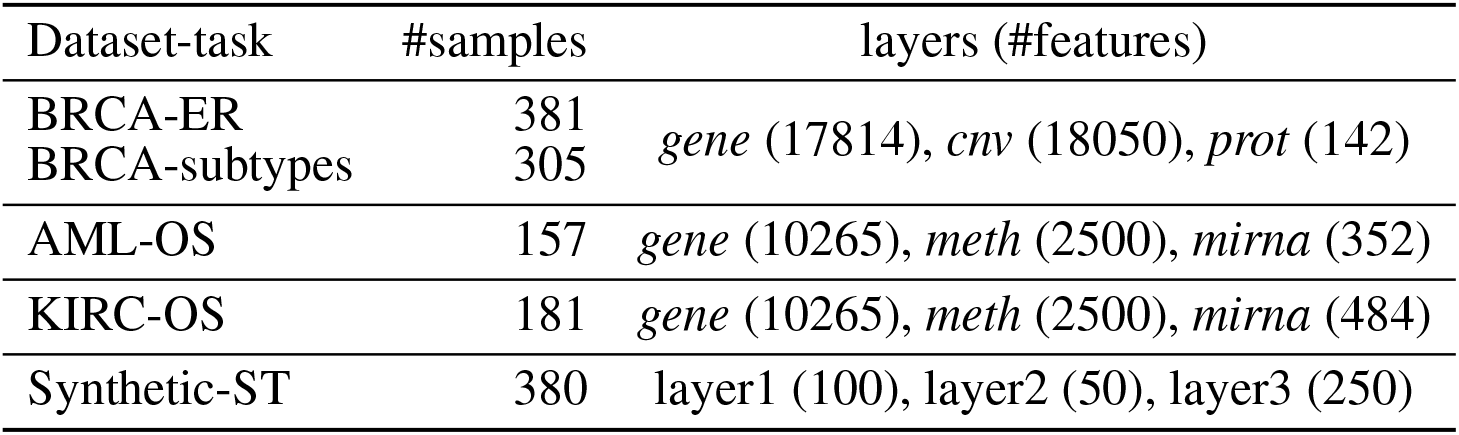
Data summary. BRCA: breast invasive carcinoma; AML: acute myeloid leukemia; KIRC: kidney renal clear cell carcinoma; *gene*: gene expression; *cnv*: copy number variants; *prot*: protein abundance; *meth*: methylation; *mirna*: microRNA expression; ER: estrogen receptor; subtypes: breast cancer subtypes; OS: overall survival; ST: synthetic target.

**Table 2.**
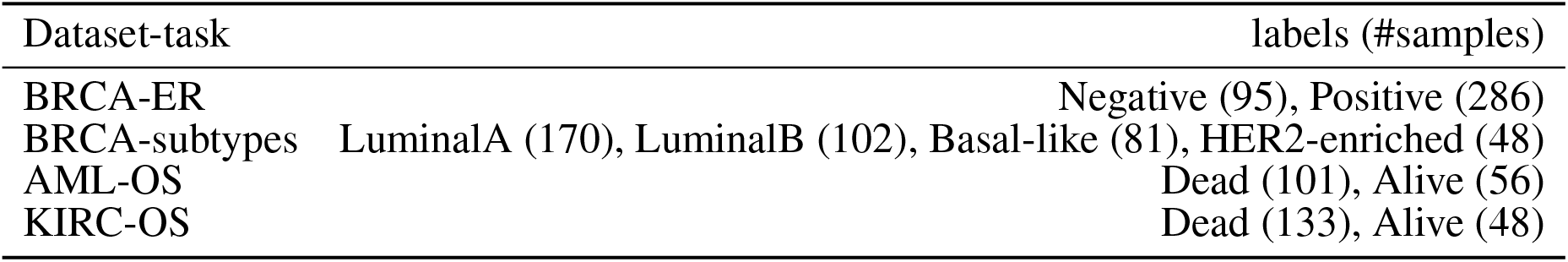
Class balance. BRCA: breast invasive carcinoma; AML: acute myeloid leukemia; KIRC: kidney renal clear cell carcinoma; ER: estrogen receptor; subtypes: breast cancer subtypes; OS: overall survival.

For AML and KIRC, gene expression is profiled using the Illumina HiSeq2000 and quantified as log2-transformed RSEM normalized counts; miRNA mature strand expression is profiled using the Illumina Genome Analyzer and quantified as reads per million miRNA mapped; and methylation is assessed by Illumina Human Methylation 450K and expressed as beta values. For BRCA, gene expression is profiled with Agilent 244K custom gene expression microarrays; protein abundance is assessed by reverse phase protein arrays; copy number profiles are measured using Affymetrix Genome-Wide Human SNP Array 6.0 platform, copy number variants are segmented by the TCGA Firehose pipeline using GISTIC2 method, and then mapped to genes.

The original data is publicly accessible on the National Cancer Institute GDC Data Portal (https://portal.gdc.cancer.gov/) and the Broad GDAC Firehose (https://gdac.broadinstitute.org/), where further details on data generation can be found. The data was retrieved in December, 2019 and January, 2020 using the *RTCGA* R library (13).

Furthermore, the INF pipeline has been tested on a synthetic dataset with 380 observations in two classes (70% class 1 and 30% class 2, defining the synthetic target ST), 3 pseudo-omics layers, and 400 features (layer 1: 100; layer 2: 50; layer 3: 250). The dataset is generated in-house using *scikit-learn*’s `make_classification` function with the arguments `shuffle=False` and `flip_y=0`. The number of informative features and the difficulty of the task were set on a per-layer basis, as summarized in Table 3.

**Table 3.** Synthetic data summary for each simulated layer. Multiplicative factor, class separation, and random state refer to the parameters `scale`, `class_sep`, and `random_state` of the `make_classification` function in *scikit-learn*.

| Layer | # features | # informative features | Multiplicative factor | Class separation | Random state |
| --- | --- | --- | --- | --- | --- |
| Layer 1 | 100 | 10 | default | 1.0 | 1 |
| Layer 2 | 50 | 5 | default | 1.2 | 2 |
| Layer 3 | 250 | 25 | 10 | 0.8 | 3 |

### 2.2 In silico workflow

The INF pipeline integrates two or more omics layers, e.g. gene expression, protein abundance, or methylation, in a machine learning framework for improved patient classification and biomarker identification in cancer. The core consists of three main components, structured as in Figure 1, managing the integration of the omics layers and their predictive modeling. A baseline integration method (juXT) is first considered by training a Random Forest (RF) (14) or a linear Support Vector Machine (LSVM) (15) classifier on juxtaposed multi-omics data, ranking features by ANOVA F-value. Secondly, the multi-omics features are integrated by Similarity Network Fusion (SNF) (5), a method that computes a sample similarity network for each data type and fuses them into one network. INF introduces a novel feature ranking scheme (rSNF) that sorts multi-omics features according to their contribution to the SNF-fused network structure. A RF or LSVM classifier is trained on the juxtaposed multi-omics data, ranking features by rSNF. A compact RF model (rSNFi) is finally trained on the juxtaposed dataset restricted on the intersection of top-ranked biomarkers from juXT and rSNF.

**Figure 1.**
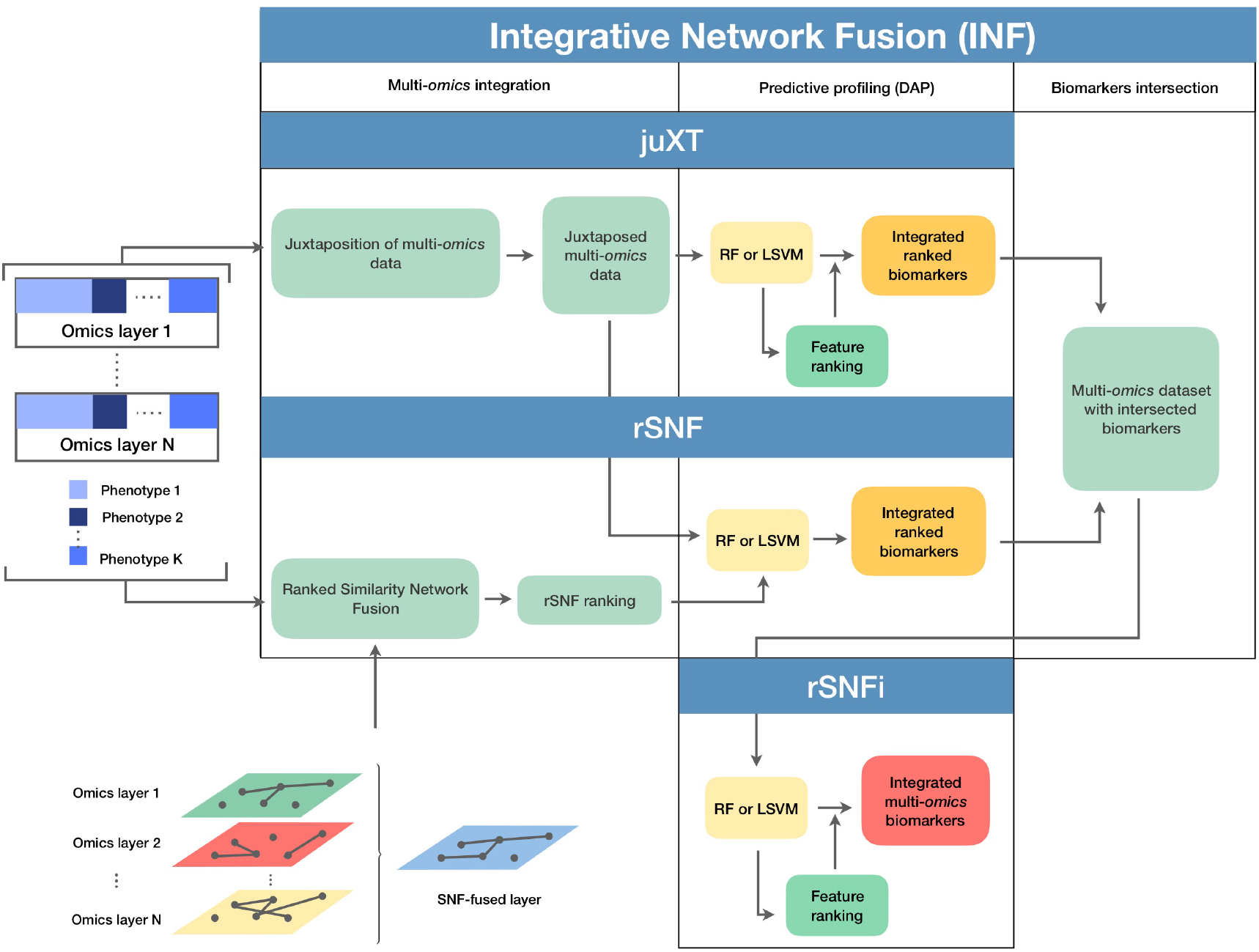
Graphical representation of the INF workflow for N *omics* datasets with K phenotypes. A first RF or LSVM classifier is trained on the juxtaposed data, ranking features by ANOVA F-value (*juXT*). The two data sets are then integrated by Similarity Network Fusion, the features are ranked by rSNF and a RF or LSVM model is developed on the juxtaposed dataset with the rSNF feature ranking (*rSNF*). Finally, a RF or LSVM classifier is trained on the juxtaposed dataset restricted to the intersection of *juXT* and *rSNF* top discriminant feature lists (*rSNFi*). The classifier is either RF or LSVM throughout the INF workflow. All the predictive models are developed within the DAP described in the methods.

### 2.3 Omics integration

In a comparative review of scientific literature, SNF (5) emerged as one of the most reliable alternatives to simple juxtaposition-based integration. SNF is a non-Bayesian network-based method that can be divided into two main steps: the first step builds a sample-similarity network for each omics dataset, where nodes represent samples and edges encode a scaled exponential Euclidean distance kernel computed on each pair of samples; the second step implements a nonlinear combination of these networks into a single similarity network through an iterative procedure. The multi-omics datasets are first converted into graphs, and for each graph two matrices are computed: a patient pairwise similarity matrix (“status matrix”), and a matrix with similarity of each patient to the K most similar patients, through K-nearest neighbors (“local affinity matrix”). At each iteration, the status matrix is updated through the local affinity matrix, generating two parallel interchanging processes. The status matrices are finally fused together into a single network. Spectral clustering is performed on the fused network, in order to identify sub-communities of samples, potentially reflecting phenotypes. The clustering performance is evaluated with respect to a ground truth, *i.e.*, the real phenotype each sample belongs to, by the Normalized Mutual Information (NMI) score. SNF integrates multiple omics datasets into a single comprehensive network in the space of samples rather than measurements (*e.g.*, gene expression values).

This work proposes multi-omics integration as an approach to identify robust biomarkers of samples phenotypes or cancer subtypes (*e.g.*, survival status vs breast cancer subtyping); consequently, it is necessary to extract measurements information from the SNF-fused network of samples. To this aim, we extended SNF by implementing *rSNF* (ranked SNF), a feature-ranking scheme based on SNF-fused network clustering. In detail, a patient network *W_i_* is built for each feature *f_i_*, based on *f_i_* alone, and spectral clustering is performed on it. Then, NMI score is computed comparing the samples clusters found inside *W_i_* with those in the fused network; the higher the score, the more similar the clustering between the fused network and *W_i_*. Thus, each feature *f_i_* is associated to a consistency score, ranking all multi-omics features with respect to their relative contribution to the whole network structure.

The entire procedure of similarity networks inference and fusion relies on two hyperparameters: *α*, the scaling variance in the scaled exponential similarity kernel used for similarity networks construction, and K, the number of nearest neighbors in sparse kernel and scaled exponential similarity kernel construction. While the original method (5) assigned fixed values to *α* and K, in this study the optimal hyperparameters are chosen among the grids *α_grid_* = {0.3, 0.35, 0.4, 0.45,…, 0.8} and *K_grid_* = {*i* ∈ ℕ, 10 ≤ *i* ≤ 30} in a 10×5-fold cross-validation schema.

### 2.4 Predictive profiling

To ensure the reproducibility of results and limit overfitting, the development of classification models is performed inside a Data Analysis Plan (DAP), following the guidelines derived by the U.S. Food and Drug Administration MAQC/SEQC studies (6, 16). Data is split in a training set (TR) and two non-overlapping test sets (TS, TS2), preserving the original proportion of patient phenotypes (classes). The TR/TS/TS2 partitions are 50%/30%/20% of the entire data set, respectively. The data splitting procedure is repeated 10 times so to obtain 10 different TR/TS/TS2 splits. Predictive models are trained and developed on TR and TS for juXT and rSNF; in the case of rSNFi, the models are trained and developed on TS and TS2 to avoid information leakage due to using the same data both for feature selection and model training (see Figure 2).

**Figure 2.**
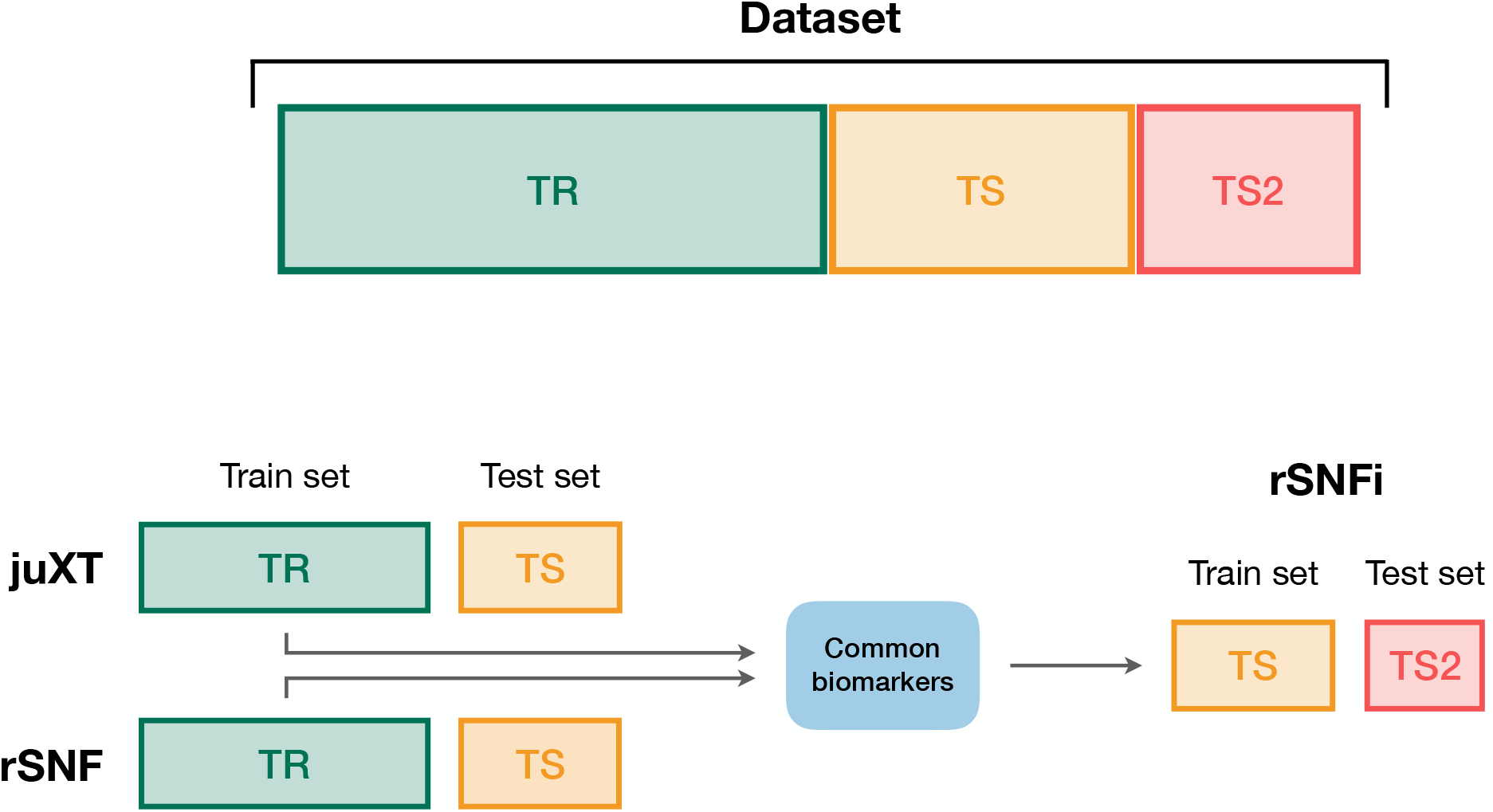
Data splitting procedure. To avoid information leakage due to the use of the same data both for feature selection and model training, we considered different train and test sets according to the integration scheme. In particular, each data set is split into three non-overlapping partitions (TR/TS/TS2), corresponding to the 50%/30%/20% of the entire data set, respectively. The TR/TS/TS2 partitions preserve the original proportion of patient phenotypes. Predictive models for juXT and rSNF are trained on TR and validated on TS, while for rSNFi the train set is TS (with features restricted to the intersected biomarkers of juXT and rSNF) and TS2 the test set.

For each split, Random Forest (RF) or linear kernel Support Vector Machine (LSVM) classifiers are trained on the training partition within a stratified 10×5-fold cross-validation (10×5-CV). The model performance is assessed in terms of average Matthews Correlation Coefficient (MCC) (17, 18), which is generally regarded as a balanced measure of accuracy and precision that can be used both in binary and multiclass problems (19, 20) and even when classes are imbalanced (21). MCC lies in [−1, 1], with 1 meaning perfect prediction, −1 inverse prediction and 0 random guess. For binary classification tasks, MCC is calculated on true and predicted labels considering true positive (TP), true negative (TN), false positive (FP) and false negative (FN) values, as in the following:

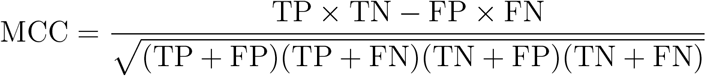

At each CV round, features are ranked either by ANOVA F-value (for juXT, rSNFi) or by the rSNF ranking (see Section 2.3) and different classification models are trained for increasing numbers of ranked features, namely 5%, 10%, 25%, 50%, 75%, and 100% of the total features. A unified list of top-ranked features is then obtained by Borda aggregation of all the ranked CV lists (22, 23). The best model is later retrained on the whole training set restricted to the features yielding the maximum MCC in CV, and validated on the test partition. A global list of top-ranked features is derived for juXT, rSNF, and rSNFi by Borda aggregation of the Borda lists of each TR/TS split (Borda of Bordas, “BoB”). The signatures for juXT, rSNF, and rSNFi are defined by the top N features of the corresponding BoB lists, with N being the median size of top features across all experiments.

In the “full” version of the DAP (*fDAP*), described above, the rSNF ranking is performed at each CV round on the training portion of the data. Since this procedure is quite demanding in terms of computational time, even if parallelized (≈ 9 feature/min), we devised an “accelerated” version of the DAP (*aDAP*), where the rSNF ranking is precomputed on the whole TR data and used as is at each CV round. We assessed the *fDAP* vs *aDAP* performance on the synthetic dataset as well as BRCA-ER and BRCA-subtypes by comparing the overall metrics and measuring the dissimilarity of the rSNF BoB of the two DAPs by the Canberra distance (22).

RF models are trained using 500 trees, measuring the quality of a split as mean decrease in the Gini impurity index (14); the regularization parameter *C* of LSVM models is tuned over the grid *C_grid_*= {10^*i*^, *i* ∈ ℕ, −2 ≤ *i* ≤ 3} within a 10× stratified Monte Carlo cross-validation (50% training/validation proportion).

To ensure that the predictive profiling procedure is not affected by selection bias, the whole INF workflow, including the rSNF procedure, is also repeated after randomly scrambling the training set labels (“random labels” mode): in the absence of systematic bias, MCC is expected to be close to the random guess value of zero.

### 2.5 Implementation

The complete INF pipeline is implemented through the workflow management tool Snakemake (24, 25), which allows automatic handling of all dependencies required to generate the INF output. The pipeline operates on N omics input files, one for each layer that should be integrated, and a single file describing the patient labels. The omics files are tab-separated text matrices with patients on the rows and features on the columns, with row and column identifiers. The label file is a single column file with patient phenotypes, with no header. This input structure, with one file per omic layer and a label file, simplifies the downstream analysis and reduces to a minimum the preprocessing burden for the end user.

The predictive profiling module, including the DAP, is written in Python 3.6 on top of NumPy (26) and scikit-learn methods (27). The ranked SNF (rSNF) procedure is implemented in R (28) leveraging the original R scripts provided by SNF authors (5), extended by a dedicated script for SNF tuning and a main script for SNF analysis and the post-SNF feature selection procedure, which is parallelized over the features for efficiency using the `foreach` R library.

All the INF code is available on the GitLab repository https://gitlab.fbk.eu/MPBA/INF.

### 2.6 Computational details

The INF computations were run on the FBK Linux high-performance computing facility KORE, on a 8-core i7 3.4 GHz Linux workstation, and on a 72-vCPU 2.7 GHz Platinum Intel Xeon 8168 Microsoft Azure cloud machine (F72s v2 series).

## 3 RESULTS

The INF workflow was run on all tasks considering 3-layer integration and all 2-layer combinations; the DAP was also run separately on all single-layer datasets in order to obtain a baseline. All results presented here refer to experiments performed with RF classifier. Experiments using LSVM were performed on BRCA-ER and KIRC-OS obtaining similar classification performances, top features and layer contributions (Supplementary Material tables *BRCA-ER_LSVM*, *KIRC-OS_LSVM*). The classifier performance for 3-layer integration is summarized in Table 4, in terms of average cross-validation MCC on the 10 training set splits (MCC_cv) with 95% Studentized bootstrap confidence intervals (CI) as (MCC_min, MCC_max), average MCC on the 10 test set splits (MCC_ts) with CI, and median number of features (Nf) yielding MCC_cv. The classifier performance on single-layer and 2-layer data is summarized in Figure 3.

**Figure 3.**
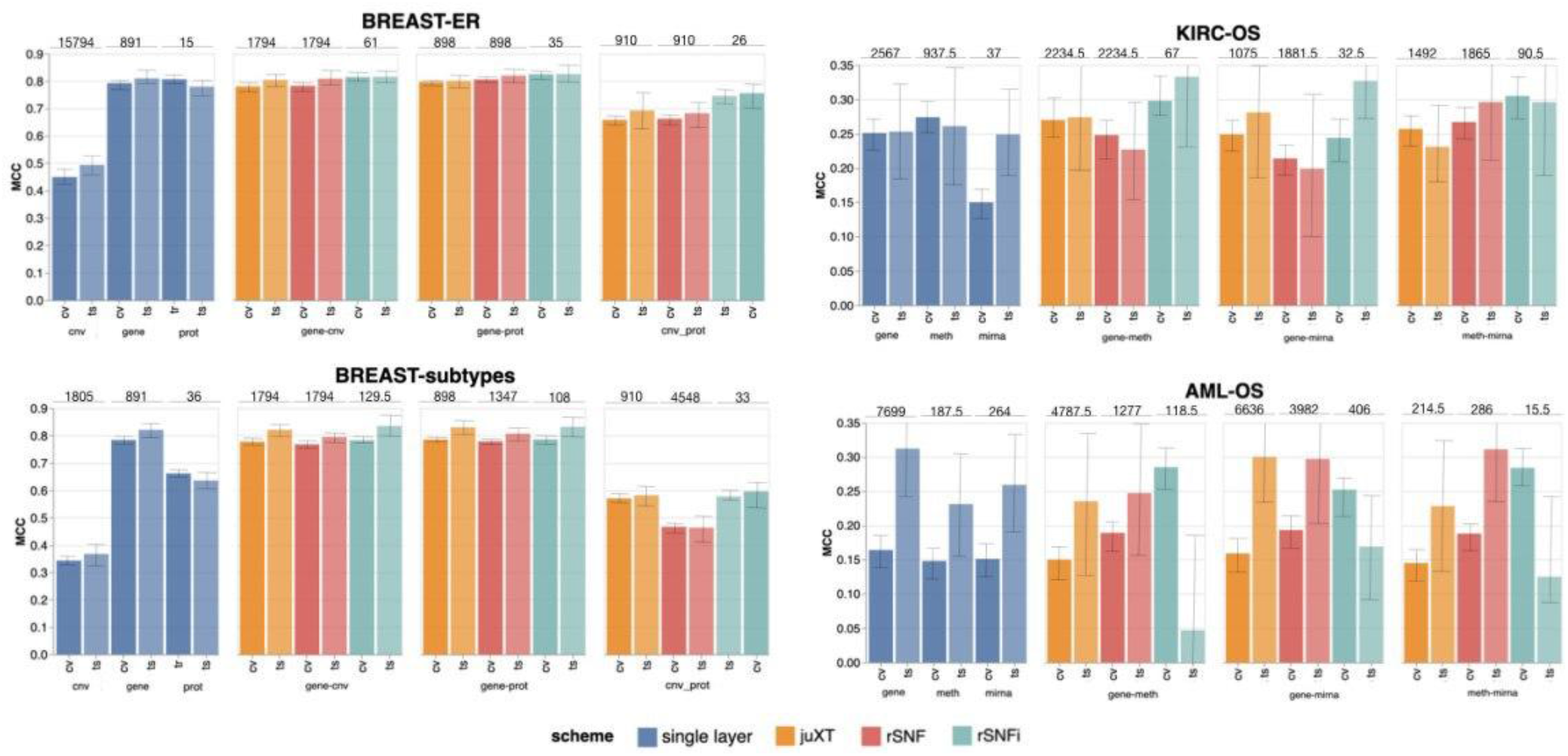
Overview of Random Forest classification performance (MCC, Matthews Correlation Coefficient) on the four tasks in cross validation (“CV”) and test (“ts”), on single-layer (blue shades) and on all two-layer combinations for juXT (orange), rSNF (red) and rSNFi (green). Bars indicate 95% confidence intervals. On top of each CV-ts pair is the median number of features leading to best CV performance.

**Table 4.** Summarized best predictive performances for each classification task using RF model and three omics layers. CI: 95% bootstrap confidence interval; MCC_cv: best average MCC in cross-validation on training set splits; MCC_ts: average MCC on validation set splits; Nf: median number of features leading to MCC_cv. Bold indicates best performance (highest MCC and smallest signature size).

| Task | Method | MCC_cv (CI) | MCC_ts (CI) | Nf |
| --- | --- | --- | --- | --- |
| BRCA-ER | juXT | 0.785 (0.776, 0.795) | 0.797 (0.778, 0.819) | 1801 |
|  | rSNF | 0.792 (0.782, 0.801) | 0.804 (0.779, 0.830) | 1801 |
|  | **rSNFi** | **0.820 (0.808, 0.831)** | **0.830 (0.803, 0.857)** | **55.5** |
| BRCA-subtypes | juXT | 0.778 (0.771, 0.785) | 0.795 (0.771, 0.817) | 1801 |
|  | rSNF | 0.769 (0.762, 0.777) | 0.811 (0.787, 0.835) | 1801 |
|  | **rSNFi** | **0.788 (0.778, 0.798)** | **0.838 (0.794, 0.879)** | **301.5** |
| KIRC-OS | juXT | 0.266 (0.243, 0.289) | 0.305 (0.229, 0.382) | 2319 |
|  | rSNF | 0.253 (0.230, 0.276) | 0.274 (0.189, 0.348) | 3313 |
|  | **rSNFi** | **0.268 (0.239, 0.298)** | **0.378 (0.288, 0.464)** | **111** |
| AML-OS | juXT | 0.141 (0.120, 0.163) | 0.223 (0.146, 0.307) | 6559 |
|  | rSNF | 0.180 (0.157, 0.202) | **0.263 (0.175, 0.366)** | 656 |
|  | **rSNFi** | **0.274 (0.245, 0.301)** | 0.176 (0.068, 0.278) | **91.5** |

A comparison between the “accelerated” flavour of the DAP (*aDAP*) and the full DAP (*fDAP*) was run on synthetic data, BRCA-ER and BRCA-subtypes data, with *aDAP* yielding similar performance metrics and top-ranked biomarker lists as *fDAP* (Supplementary Material tables *Synthetic_RF*, *BRCA_RF_fDAP*, *canberra_distances*), while being ≈ 30× faster (for BRCA-ER, approx. 2h vs 64h, or 300 features/min vs 9 features/min). All the results presented here were thus obtained using *aDAP*. Moreover, the INF workflow running in “random labels” mode achieved an average cross-validation MCC ≈ 0, as expected by a procedure unaffected by systematic bias.

Overall, integrating multiple omics layers with INF yields better or comparable classification performance than using only features from a single layer or naïve omics juxtaposition, at the same time with much more compact signature sizes. On 3-layer BRCA-subtypes and 2- or 3-layer KIRC-OS, INF outperforms the single layers, as well as juXT and rSNF (Figure 3, Table 4). On 2-layer BRCA-subtypes, INF performance on *gene*-*cnv* and *gene*-*prot* is comparable to the best-performing single-layer data (*gene*) and superior to *cnv* and *prot* single layers, while INF on *cnv*-*prot* only improves over the *cnv* single layer. On the BRCA-ER task, the performance with INF integration of 2 or 3 layers is still better than using single layers, nevertheless to a smaller extent, except for *cnv*-*prot* integration which performs better than *cnv* alone but slightly worse than *gene* and *prot* single layers. On the more difficult AML-OS task, INF has better performance over both rSNF and juXT on *gene*-*mirna* and *meth*-*mirna* integration, still improving over single-layer performance both in terms of MCC and reduced signature sizes.

### One or multi-omics layers vs juXT/rSNF/rSNFi

For BRCA-ER, three-layer INF (rSNFi) integration performs better than either rSNF or juXT (MCC test 0.830 vs 0.804, 0.797 for rSNF and juXT, respectively). All two-layer INF integrations perform similarly to, or better than, the corresponding rSNF and juXT integrations, in particular for *cnv*-*prot* integration (MCC test 0.746 vs 0.682, 0.692 resp. for rSNF and juXT).

On BRCA-subtypes, the 3-layer INF integration performs better than either rSNF or juXT (MCC test 0.838 vs 0.811, 0.795 resp. for rSNF and juXT), nevertheless without improving over the *gene* single-layer performance (MCC test 0.821). However, the INF median signature size is only 301.5, compared to 1801 for rSNF and juXT, and 891 for the *gene* layer alone. All two-layer INF integrations yield better performance than their corresponding juXT or rSNF integrations.

Omics integration is particularly effective for KIRC-OS, as all 2- and 3-layer INF integrations outperform juXT, rSNF, and each of the single-layer classifiers. In fact, 3-layer rSNFi achieves MCC test 0.378 vs 0.274, 0.305 (resp. for juXT, rSNF), 0.296, 0.327, 0.333 (resp. rSNFi *meth*-*mirna*, *gene*-*mirna*, *gene*-*meth*), and 0.253, 0.261, 0.249 (resp. *gene*, *meth*, *mirna*).

For AML-OS, INF feature sets are always more compact than either juXT or rSNF, with three-layer integration giving better MCC than any of the INF two-layer integrations (MCC test 0.176 vs 0.125, 0.169, 0.047, respectively three-layer vs *meth*-*mirna*, *gene*-*mirna*, *gene*-*meth*). Moreover, cross-validation MCCs corresponding to INF integration are better than any single layer MCC as well as rSNF and juXT.

### Characterization of the signatures identified by INF

For all tasks, INF signatures are markedly more compact with respect to both juXT and rSNF. With 91.5 vs 6559 (1.4%) median features (rSNFi vs juXT), the largest reduction in size occurs for AML-OS 3-layer integration, while the least reduction is observed for BRCA-subtypes task, with 301.5 vs 1801 (16.7%) median features (rSNFi vs juXT).

In terms of contributions from the omics datasets being integrated, the *gene* layer generally provides the largest number of features to the signatures identified by the INF workflow. In particular for the BRCA dataset, in both ER and subtypes tasks, the *gene* layer contributes over 95% of the top features for juXT and rSNFi, with rSNF signatures being slightly more balanced (*prot* contribution remains marginal, while *cnv* provides 28.3% and 17.7% of the top features in ER and subtypes tasks respectively). In AML-OS experiments, the layer contributing the most is still *gene*, accounting for ca. 78%, 73% and 81% of the top feature sets for RF juXT, rSNF and rSNFi experiments, respectively. In KIRC-OS experiments, *gene* is the layer contributing the most to the top juXT and rSNF feature sets, while *meth* is the major contributor for rSNFi. The percentage of features from each omic layer contributing to the top signatures for juXT, rSNF and rSNFi 3-layer integrations are reported in Supplementary Material table *layer contribution*. The RF rSNFi signatures for all tasks are available in Supplementary Material tables *BRCA-ER_RF_rSNFi*, *BRCA-subtypes_RF_rSNFi*, *AML-OS_RF_rSNFi* and *KIRC-OS_RF_rSNFi*.

Even though a systematic biological interpretation of the signatures identified is beyond the scope of this work, to ascertain the reliability of our results we compared them with published data. The top features in the BRCA-ER rSNFi signature include multiple genes known to be associated with breast carcinoma progression and outcome such as AGR3, B3GNT and MLPH (29, 30, 31). In addition we find the estrogen receptor gene (ESR1 from the *gene* and ER-alpha from the *prot* layer) and the transcription factor GATA3 (from both *gene* and *prot* layers) (32). Both the BRCA-ER and BRCA-subtypes signatures include genes previously identified as novel biomarkers for intrinsic breast carcinoma subtype prediction (33). Interestingly there is only partial overlap between the top features identified in BRCA ER vs subtypes tasks. Considering AML-OS task, it is noteworthy to mention that the top feature identified has been recently reported as a potential biomarker predicting overall survival in a subset of AML patients (34).

Within the *mirna* features of the AML-OS signature, MIR-203 expression was recently found to be associated with AML patient survival (35); MIR-100 is highly expressed in AML and was found to regulate cell differentiation and survival (36); high expression of miR-504-3p was reported to be associated with favorable AML prognosis (37). Given that the rSNFi signature identified in the KIRC-OS task contains a large percentage of methylation data (86.5%), its direct interpretation is more difficult. It is however interesting to observe that all the 15 *gene* features in the signature are identified as prognostic markers for renal carcinoma according to the Human Protein Atlas (38).

### Unsupervised analysis

The features selected by juXT, rSNF and rSNFi are projected on a bi-dimensional space using the UMAP unsupervised multidimensional projection method (39, 40). Here we show an example on the BRCA-subtypes 3-layer dataset, with a UMAP projection of the features selected by juXT (Figure 4(a)) compared to the UMAP projection of the INF signature (Figure 4(b)) for one of the 10 data splits (the UMAP plots for the remaining 9 splits are in Supplementary Material figures S1, S2). Colors represent cancer subtypes and shapes represent training/test partitions. Using the 1801 juXT features, cancer subtypes are roughly clustered, with HER2-enriched and Luminal B being more dispersed (Figure 4(a). The clusters appear to be more sharply defined in the projection of the 302-feature INF signature: in particular, Basal-like patients form a distinct cluster, while Luminal A, Luminal B and HER2-enriched patient clusters are close to each other, slightly overlapping yet hinting to a trajectory pattern (Figure 4(b)). The HER2/luminal cluster contains two patients classified as basal-like subtype, consistently with the findings of Koh and colleagues (41).

**Figure 4.**
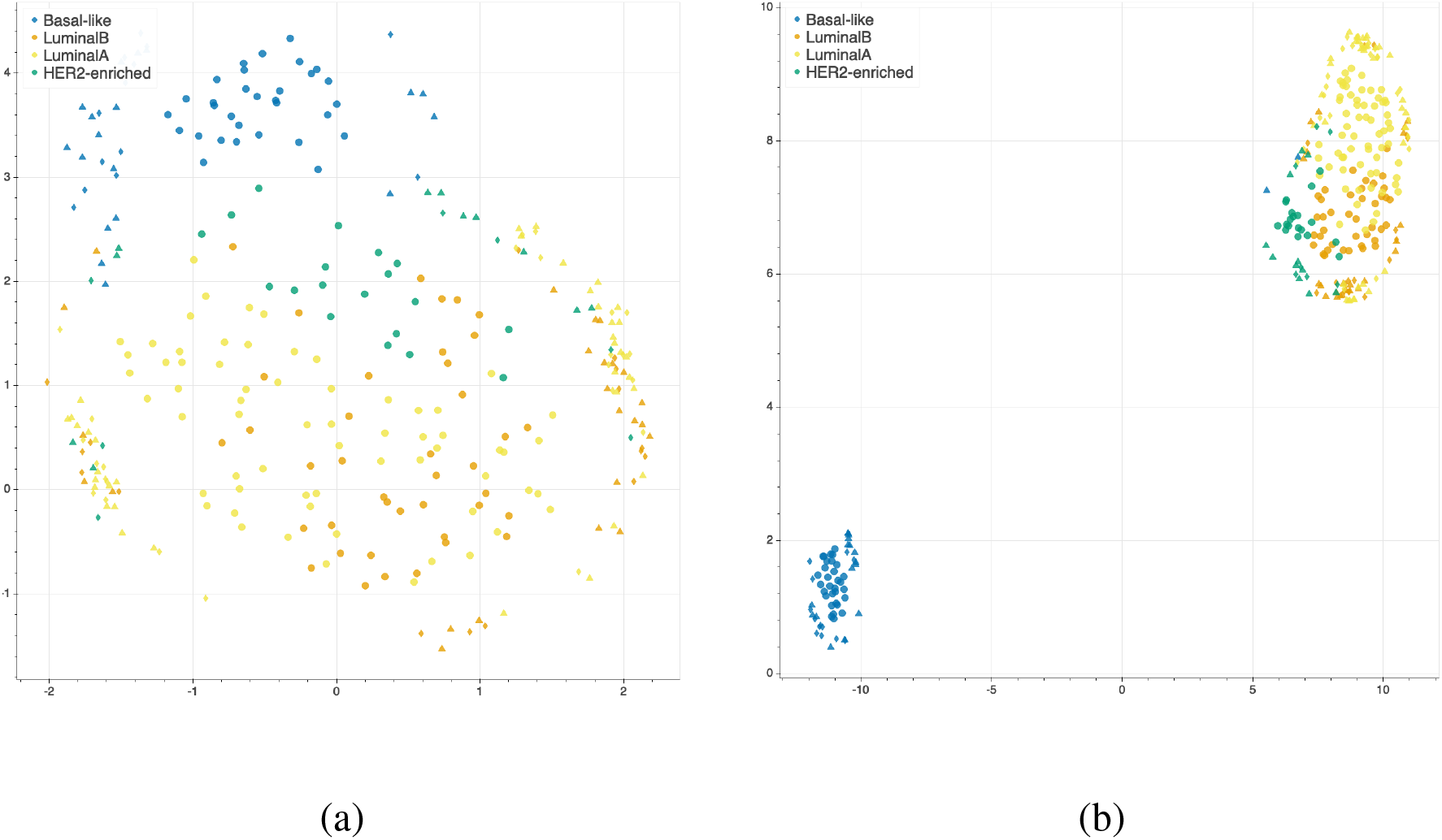
UMAP projection on the BRCA-subtypes task with 3-layer juxtaposed data (a) and restricted to the rSNFi signature (b). Circle: TR set; triangle: TS set; diamond: TS2 set.

## 4 DISCUSSION

### 4.1 Background and related work

Ritchie and colleagues (42) defined omics data integration as the combination of multiple omics datasets that can be used for the development of models to predict complex traits or phenotypes. The problem of data integration in computational biology is far from having a consolidated and shared solution. Many long-standing obstacles are still far from being overcome, and the increasing availability of data (e.g., TCGA, (43)) and computational tools (see for instance (44, 45, 46, 47, 48) and https://github.com/mikelove/awesome-multi-omics), also interactive (e.g. (49)), is raising new issues that need to be addressed. In fact, not only are existing datasets still lacking standardization protocols to deal with their complexity and heterogeneity, but also the reliability, reproducibility and interpretability of new computational methods are emerging as urgent and relevant questions (50). Moreover, modern technologies allow the rapid extraction of high-dimensional, high-throughput features from different sources (e.g. gene expression, DNA sequencing, metabolomics, or high-resolution images), which in turn require collaboration between biologists, computer scientists, physicians and other experts. The lack of common methodologies and terminologies can transform this synergy into a further level of complexity in the process of data integration (51). As observed in (52, 53), specific technological limits, noise levels and variability ranges affect the different omics, and thus confounding the underlying biological signals, yielding that really integrative analysis is still very rare, while different methods often discover different kinds of patterns, as evidenced by the lack of consistency in the published results, although efforts in this direction have started appearing (54, 55).

Indeed, the underlying hypothesis of multi-omics integration is that different omics data can provide complementary information (53) (although sometimes redundant (9)), and thus a broader insight with respect to single-layer analysis, for a better understanding of disease mechanisms (56). This assumption has been confirmed by multiple studies on diverse diseases, such as cardiovascular disease (57), diabetes (58), liver disease (59), or mitochondrial diseases (60), and also longitudinally (61), suggesting that the more complex the disease the more advantageous the integration. As the co-occurence of multiple causes and correlated events is a well-known characteristic of tumorigenesis and cancer development, the integration of data generated from multiple sources can thus be particularly useful for the identification of cancer hallmarks (62, 63, 64, 65).

Many computational strategies have been introduced that combine multiple types of data to identify novel biomarkers and thus to predict a phenotype of interest or drive the development of intervention protocols. Given the heterogeneity of data and tasks, these techniques deal with the data integration at different levels of the learning process: (i) by concatenating the features before fitting a model (early-integration), (ii) by incorporating the integration step into the model training (intermediate-integration), or (iii) by combining the outputs of distinct models for the final prediction (late-integration) (66, 67).

In the early-integration approach, also known as juxtaposition-based, the multi-omics datasets are first concatenated into one matrix. To deal with the high-dimensionality of the joint dataset, these methods generally adopt matrix factorization (68, 53, 55, 52), statistical (46, 69, 70, 59, 57, 44, 71, 72, 73, 55), and machine learning tools (74, 73, 55). Although the dimensionality reduction procedure is necessary and may improve the predictive performance, it can also cause the loss of key information (66). Moreover, biomarkers identified purely on a computational statistics rationale from meta-omics features often lack biological plausibility (75).

In order to maximize the contribution of the single-omics layer, the late-integration methods first model each dataset individually, and then merge or average the results; they are also known as model-driven (76, 67). Although these techniques avoid the pre-selection of the features, they do not leverage the hidden correlations between the data, posing again the risk of signal loss (77, 75).

The intermediate-integration strategies aim at developing a joint model that accounts for the correlation between the omics layers, to boost their combined predictive power (78). Among these methods, the network-based models refer to the reconstruction of a graph representing the complex biological interactions (79, 73), known or predicted, between the variables to discover novel informative relationships (80). They have successfully been applied in cancer research for the identification of pan-cancer drug targets (81), the detection of subtype-specific pathways (82, 78) and of genetic aberrations (83), or the stratification of cancer patients (84, 85, 86). In particular, Koh and colleagues (41) predicted breast cancer subtypes by applying a modified shrunken centroid method in the development of their network-based tool, iOmicsPASS. Further, breast cancer datasets in TGCA represent a benchmark for integrative models (87, 88, 89), as well as AML (90).

More recently, the success of deep learning algorithms in various bioinformatics fields (91) prompted the adoption of deep neural network for omics-integration in precision oncology. Autoencoders and convolutional neural networks have been effectively trained for the prediction of prognostic outcomes (92, 9), response to chemotherapeutic drugs (47), and gene targeting (93), by adopting either an early-integration (9, 93) or a late-integration (92, 47). Although deep learning models hold the potential to include image-derived features in the integration workflow, they suffer from interpretability and generalization issues (94).

Although it is clear that no single method is consistently preferable, and that most of the proposed approaches are task and/or data dependent (75), the complexity of tumor analysis suggests that network-based approaches are needed (82, 95).

In this context, it is clear that omics-integration is one of the most promising and demanding challenge of the modern bioinformatics, and that there is an urgent need to prove the reproducibility, interpretability, and generalization capability of the proposed methods (80, 96).

### 4.2 Integrative Network Fusion

We present the INF framework for the characterization of cancer patient phenotypes by integrated multi-omics signatures, combining an improved version of a state-of-the-art integration technique (5) with predictive models developed inside a Data Analysis Plan (6) for machine learning. The framework is applied to TCGA data to predict clinically relevant patient phenotypes such as the overall survival or cancer subtypes.

The simplest approach for multi-omics data integration consists in juxtaposition of normalized measurements into one joint matrix, followed by the development of a predictive model. Juxtaposition-based integration is considered as a baseline technique, since it is the most naïve approach to combine two datasets; moreover, it enables to identify multi-omics signatures by borrowing discriminatory strength from information derived by all datasets. Juxtaposition further dilutes the already possible low signal-to-noise ratio in each data type, affecting the understanding of the biological interactions at the different omics levels.

Conversely, our INF method for omics data integration is an improvement of the popular Similarity Network Fusion (SNF) approach (5), which has inspired several studies in the scientific literature, specifically in cancer genomics (97, 98, 99, 100, 74, 82, 101). SNF maximizes the shared or correlated information between multiple datasets by combining data through inference of a joint network-based model, accounting for how informative each data type is to the observed similarity between samples.

Two innovative solutions have been implemented in this study: (i) we devised a SNF-based procedure to rank variables according to their importance in clustering samples with similar phenotypes; and (ii) predictive models were developed exploiting the SNF-ranked variables, inside a rigorous Data Analysis Plan which ensures reproducibility (6, 16).

The performance of INF was assessed both in terms of statistical properties as well as biological interest. Concerning the statistical aspect, INF was compared with predictive models developed on the juxtaposed datasets (juXT technique), as well as on the single-layer datasets. With INF, smaller signature sizes were systematically derived to achieve comparable or even better performance both in cross-validation and in test. This is an added value for INF, as biological validation of biomarkers can definitely benefit from signatures of small size in terms of both costs and required time. This main achievement is mainly due to the novel rSNF ranking, which increases the signal-to-noise ratio from the combined layers by prioritizing the most discriminant biomarkers in terms of network mutual information. rSNF exploits two main SNF advantages: integration of heterogeneous data and clustering of sample networks. The main peculiarity of the SNF integrative procedure is its robustness to noise (5), because weak similarities among samples (low-weight edges) disappear, except for low-weight edges supported by all networks, which are conserved depending on how tightly connected their neighborhoods are across networks. Moreover, the rSNFi step further increases the signal-to-noise ratio by training a predictive classifier on multi-omics juxtaposed data restricted to the top-ranked biomarkers shared by juXT and rSNF models. The resulting signatures are compact in size (up to 99% reduction w.r.t. juXT) while allowing predictive models to achieve equal or better performance compared to naïve juxtaposition or the single layers alone. While a comprehensive evaluation of the biological meaning of the signatures identified through the INF framework is beyond the scope of this work, we assessed their general validity with a thorough literature search. Our investigation shows that the signatures identified through the INF framework include biological markers that are relevant in the tasks under analysis and are consistent with previously published data. Further, as in (9), the largest contribution in the biomarkers’ lists is provided by gene expression, while epigenomics, proteomics and miRNA transcriptomics play a minor role.

A fair comparison of INF results with other integration methods is currently unfeasible due to the number and variety of computational pipelines with dissimilar datasets, preprocessing methods, data analysis plans, and performance metrics.

This work is based on the original R implementation of the SNF algorithm (5). However, we are aware that Open Source implementations exist in other programming languages, in particular *snfpy* for Python (102). In a future release of the INF workflow, we plan to migrate the SNF-related parts to *snfpy* or a similar Python-based implementation, in order to drop the dependency on R and to potentially improve the overall performance.

In its current version, the INF framework supports the integration of two or more one-dimensional omics layers. As part of our future effort we will add support for the integration of medical imaging layers, for example leveraging the extraction of histopathological features from whole slide images by deep learning (10) or using radiomics or deep features from radiological images (11). In both cases, further issues will emerge from the interactions between the omics and the non-omics data, needing particular care in the integration (12).

## AUTHOR CONTRIBUTIONS

Conceptualization: CA, LT, GJ; methodology: MC, NB, AM, AZ, LT, CA, GJ; interpretation: MF; coordination: GJ; writing: MC, NB, AM, MF, GJ, CF.

## ACKNOWLEDGMENTS

The authors wish to thank Dr. Valerio Maggio for helpful discussions on aspects of the machine learning workflow and for paper proofreading.

## SUPPLEMENTAL DATA

Supplementary file 1 contains the additional figures S1, S2; Supplementary file 2 contains the additional tables referenced in the main text.

## DATA AVAILABILITY STATEMENT

The original datasets analyzed in this study can be found on the National Cancer Institute GDC Data Portal (https://portal.gdc.cancer.gov/).

## Footnotes

1 INF source code is publicly available on the GitLab repository https://gitlab.fbk.eu/MPBA/INF, while data is archived at http://dx.doi.org/10.6084/m9.figshare.12052995.v1

2 European Call Multi-omics for genotype-phenotype associations (RIA) https://ec.europa.eu/info/funding-tenders/opportunities/portal/screen/opportunities/topic-details/biotec-07-2020

