## Supplemental Material 1 for "Integrative Network Fusion: a multi-omics approach in molecular profiling"

Cold Spring Harbor Laboratory
bioRxiv
THE PREPRINT SERVER FOR BIOLOGY

### Supplementary Material

#### 1 SUPPLEMENTARY FIGURES

**Figure S1.** UMAP projections on the BRCA-subtypes task with 3-layer juxtaposed data. Each subplot represents the projection of the TR/TS/TS2 partition for the remaining 9 splits not reported in the main text. Circle: TR set; triangle: TS set; diamond: TS2 set.

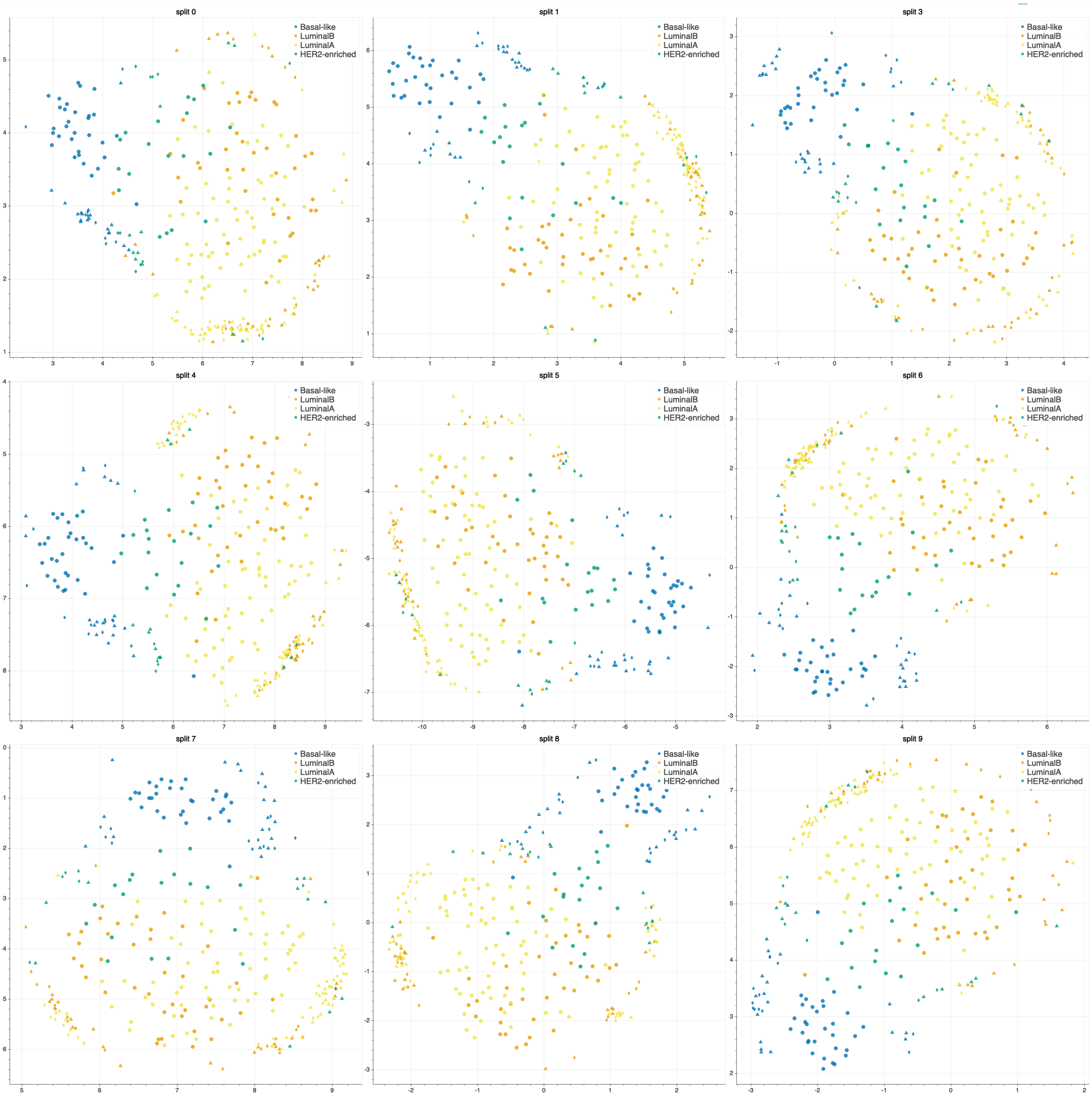

**Figure S2.** UMAP projections on the BRCA-subtypes task with 3-layer juxtaposed data restricted to the INF signature. Each subplot represents the projection of the TR/TS/TS2 partition for the remaining 9 splits not reported in the main text. Circle: TR set; triangle: TS set; diamond: TS2 set.

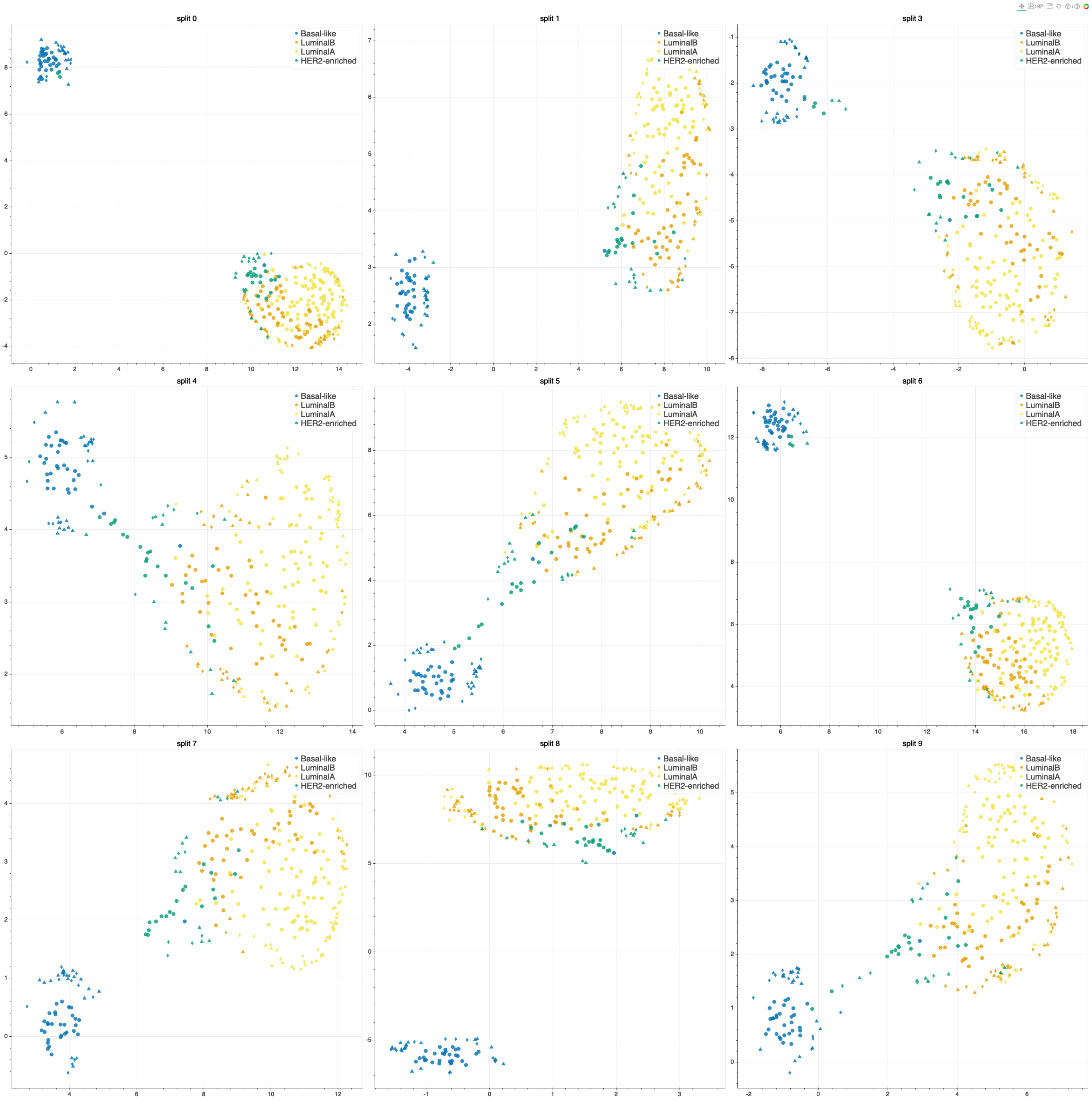
